# From genomes to phenotypes: Traitar, the microbial trait analyzer

**DOI:** 10.1101/043315

**Authors:** Aaron Weimann, Kyra Mooren, Jeremy Frank, Phillip B. Pope, Andreas Bremges, Alice C. McHardy

## Abstract

The number of sequenced genomes is growing exponentially, profoundly shifting the bottleneck from data generation to genome interpretation. Traits are often used to characterize and distinguish bacteria, and are likely a driving factor in microbial community composition, yet little is known about the traits of most microbes. We describe Traitar, the microbial trait analyzer, which is a fully automated software package for deriving phenotypes from the genome sequence. Traitar provides phenotype classifiers to predict 67 traits related to the use of various substrates as carbon and energy sources, oxygen requirement, morphology, antibiotic susceptibility, proteolysis and enzymatic activities. Furthermore, it suggests protein families associated with the presence of particular phenotypes. Our method uses L1-regularized L2-loss support vector machines for phenotype assignments based on phyletic patterns of protein families and their evolutionary histories across a diverse set of microbial species. We demonstrate reliable phenotype assignment for Traitar to bacterial genomes from 572 species of 8 phyla, also based on incomplete single-cell genomes and simulated draft genomes. We also showcase its application in metagenomics by verifying and complementing a manual metabolic reconstruction of two novel Clostridiales species based on draft genomes recovered from commercial biogas reactors. Traitar is available at https://github.com/hzi-bifo/traitar.

## Introduction

Microbes are often characterized and distinguished by their traits, for instance, in *Bergey’s Manual of Systematic Bacteriology* (Goodfellow et al., 2012). A trait or phenotype can vary in complexity; for example, it can refer to the degradation of a specific substrate or the activity of an enzyme inferred in a lab assay, the respiratory mode of an organism, the reaction to Gram staining or antibiotic resistances. Traits are also likely driving factor for microbial community composition (Martiny et al., 2015). Microbial community members with varying metabolic capabilities can aid in waste water treatment, bioremediation of soils and promotion of plant growth (Bai et al., 2015; Narihiro and Sekiguchi, 2007; Olapade and Ronk, 2015); in the cow rumen microbiota, bacterial cellulose degraders influence the ability to process plant biomass material (Hess et al., 2011). In the Tammar wallaby foregut microbiome, the dominant bacterial species is implicated in the lower methane emissions produced by wallaby compared to ruminants (Pope et al., 2011).

In addition to the exponential growth of available sequenced microbial genome isolates, metagenome and single cell genome sequencing further contributes to the increasing number of available genomes. For the recovery of genomes from metagenomes (GFMs), computational methods based on e.g. differential read coverage and *k*-mer usage were developed (Alneberg et al., 2014; Cleary et al., 2015; Gregor et al., 2016; Imelfort et al., 2014; Kang et al., 2015; Nielsen et al., 2014), which allow to recover genomes without the need to obtain microbial isolates in pure cultures (Brown et al., 2015; Hess et al., 2011). In addition, single-cell genomics provides another culture-independent analysis technique and also allows, although often fragmented, genome recovery for less abundant taxa in microbial communities (Lasken and McLean, 2014; Rinke et al., 2013). Together, these developments profoundly shift the analytical bottleneck from data generation to interpretation.

The genotype–phenotype relationships for some microbial traits have been well studied. For instance, bacterial motility is attributed to the proteins of the flagellar apparatus (Macnab, 2003). We have recently shown that delineating such relationships from microbial genomes and accompanying phenotype information with statistical learning methods enables the accurate prediction of the plant biomass degradation phenotype and the *de novo* discovery of both known and novel protein families that are relevant for the realization of the plant biomass degradation phenotype (Konietzny et al., 2014; Weimann et al., 2013). However, a fully automated software framework for prediction of a broad range of traits from only the genome sequence is currently missing. Additionally, horizontal gene transfer, a common phenomenon across bacterial genomes, has not been utilized to improve trait prediction so far. Traits with their causative genes may be transferred from one bacterium to the other (Ochman et al., 2000; Pal et al., 2005) (e.g. for antibiotic resistances (Martinez, 2008)) and the vertically transferred part of a bacterial genome might be unrelated to the traits under investigation (Barker and Pagel, 2005; Harvey and Pagel, 1991; Martiny et al., 2015).

Here we present Traitar, the microbial trait analyzer: an easy-to-use, fully automated software framework for the accurate prediction of currently 67 phenotypes directly from the genome sequence (Figure 1).

**Figure 1:**
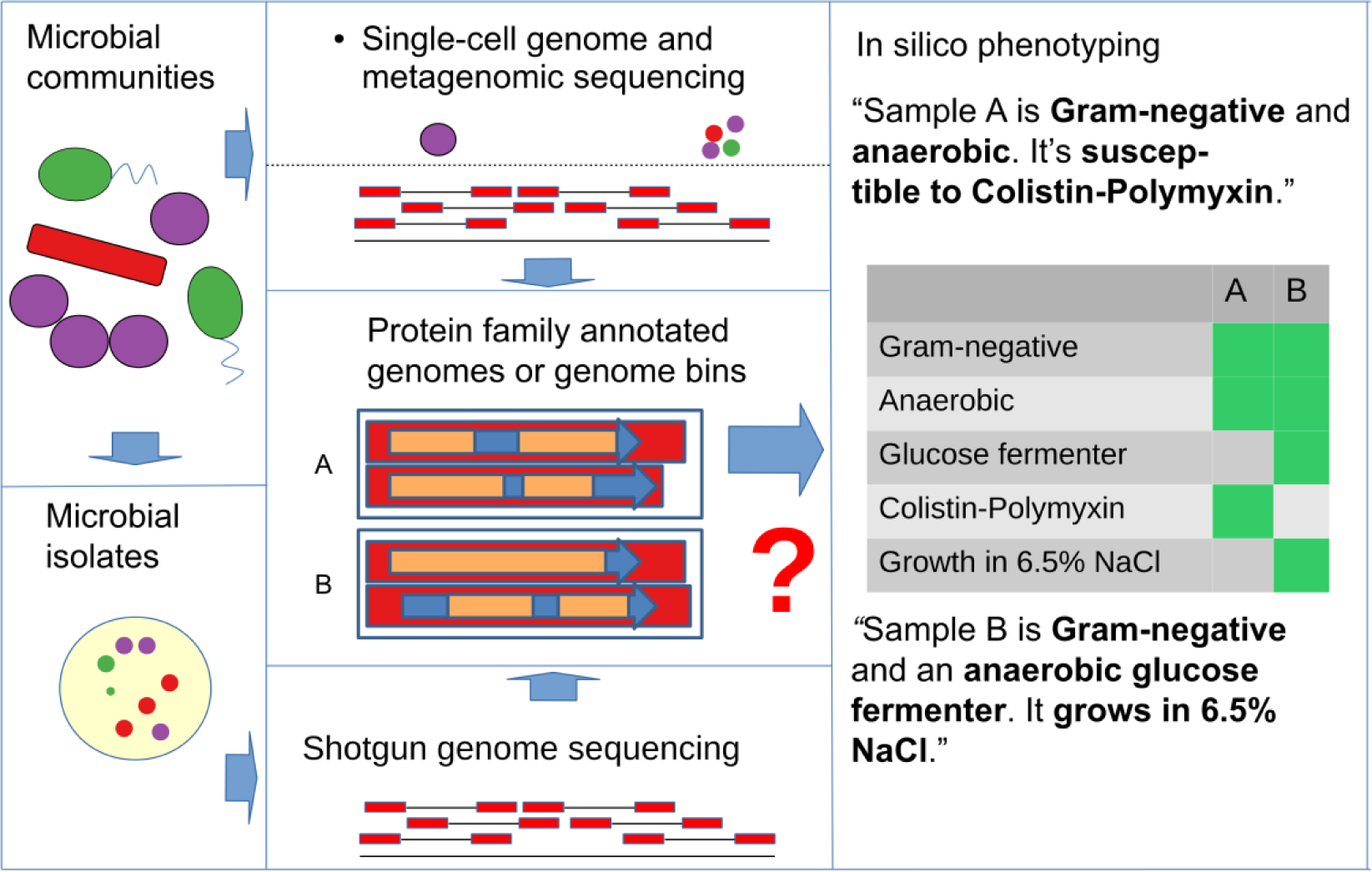
Traitar can be used to phenotype microbial community members based on genomes recovered from single-cell sequencing or (metagenomic) environmental shotgun sequencing data or of microbial isolates. Traitar provides classification models based on protein family annotation for a wide variety of different phenotypes related to the use of various substrates as source of carbon and energy for growth, oxygen requirement, morphology, antibiotic susceptibility and enzymatic activity.

We used phenotype data from the microbiology section of the Global Infectious Disease and Epidemiology Network (GIDEON) – a resource dedicated to the diagnosis, treatment and teaching of infectious diseases and microbiology (Berger, 2005) – for training phenotype classification models on the protein family annotation of a large number of sequenced genomes of microbial isolates (predominantly bacterial pathogens). We investigated the effect of incorporating ancestral protein family gain and losses into the model inference on classification performance, to allow consideration of horizontal gene transfer events in inference of phenotype-related protein families and phenotype classification. We rigorously tested the performance of our software in cross-validation experiments, on further test data sets and for different taxonomic ranks. To test Traitar’s applicability beyond the bacteria represented in GIDEON, we subsequently applied it to several hundred bacteria described in Bergey’s systematic bacteriology (Goodfellow et al., 2012). We used Traitar to phenotype bacterial single amplified genomes (SAGs) and simulated incomplete genomes to investigate its potential for phenotyping microbial samples with incomplete genome sequences. We characterized two novel Clostridiales species of a biogas reactor community with Traitar, based on their genomes recovered with metagenomics. This verified and complemented a manual metabolic reconstruction. As Traitar furthermore suggests protein families associated with the presence of a particular phenotype, we discuss the protein families Traitar identified for several phenotypes, namely for ‘Motility’, ‘Nitrate to nitrite’ conversion and ‘L-arabinose’ fermentation.

Traitar is implemented in Python 2.7. It is freely available under the open-source GPL 3.0 license at https://github.com/hzi-bifo/traitar and as a Docker container at https://hub.docker.com/r/aweimann/traitar. A Traitar web service can be accessed at https://research.bifo.helmholtz-hzi.de/traitar.

## Results

### The Traitar software

We begin with a description of the Traitar software and phenotype classifiers. Traitar predicts the presence or absence of a phenotype, i.e. assigns a phenotype label, for 67 microbial traits to every input sequence sample (Table 1, Supplementary Table 1). For each of these traits, Traitar furthermore suggests candidate protein families associated with its realization, which can be subject of experimental follow-up studies.

For phenotype prediction, Traitar uses one of two different classification models. We trained the first classifier – the phypat classifier – on the protein and phenotype presence & absence labels from 234 bacterial species (Methods – Phenotype models). The second classifier – the phypat+PGL classifier – was trained using the same data and additionally information on evolutionary protein family and phenotype gains and losses. The latter were determined using maximum likelihood inference of their ancestral character states on the species phylogeny (Methods – Ancestral protein family and phenotype gains and losses).

The input to Traitar is either a nucleotide sequence FASTA file for every sample, which is run through gene prediction software, or a protein sequence FASTA file. Traitar then annotates the proteins with protein families. Subsequently, it predicts the presence or absence of each of the 67 traits for every input sequence. Note that Traitar doesn’t require a phylogenetic tree for the input samples.

Finally, it associates the predicted phenotypes with the protein families that contributed to these predictions (Figure 2). A parallel execution of Traitar is supported by GNU parallel (Tange, 2011). The Traitar annotation procedure and the training of the phenotype models are described in more detail below (Methods – Traitar software).

**Table 1:**
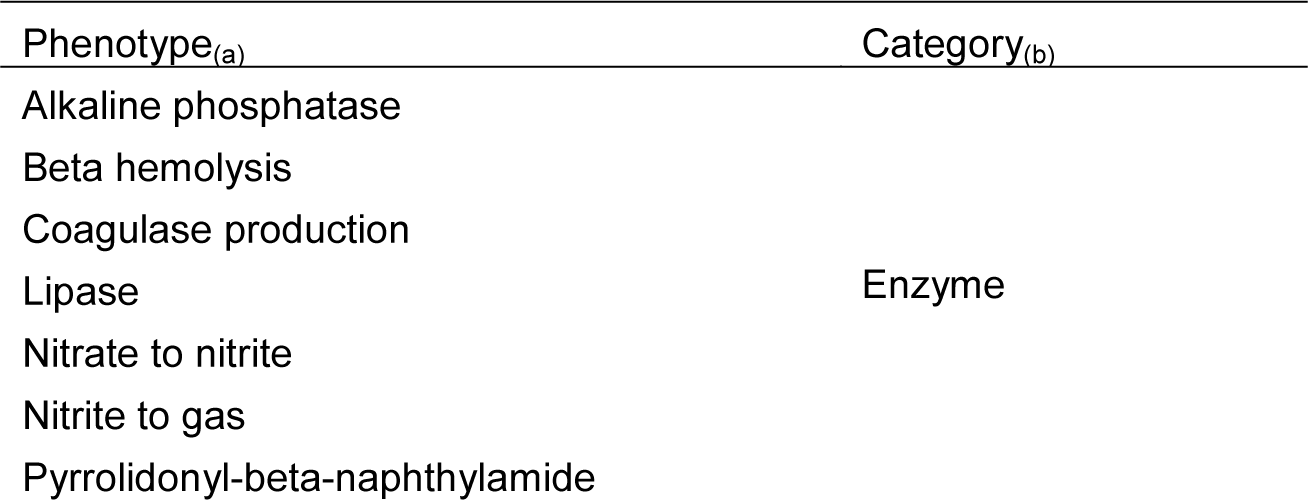
The 67 traits available in Traitar for phenotyping. We grouped each of these phenotypes into a microbiological or biochemical category.

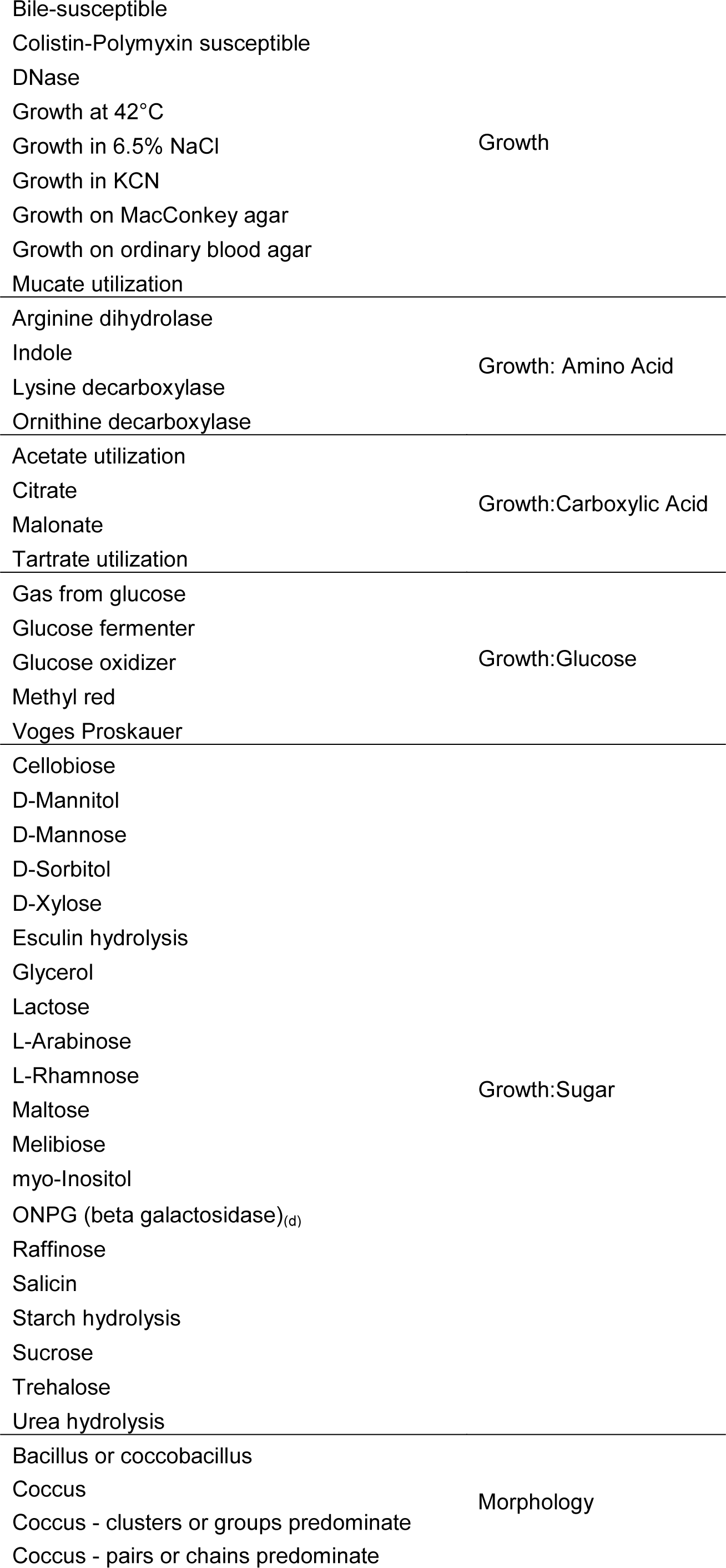

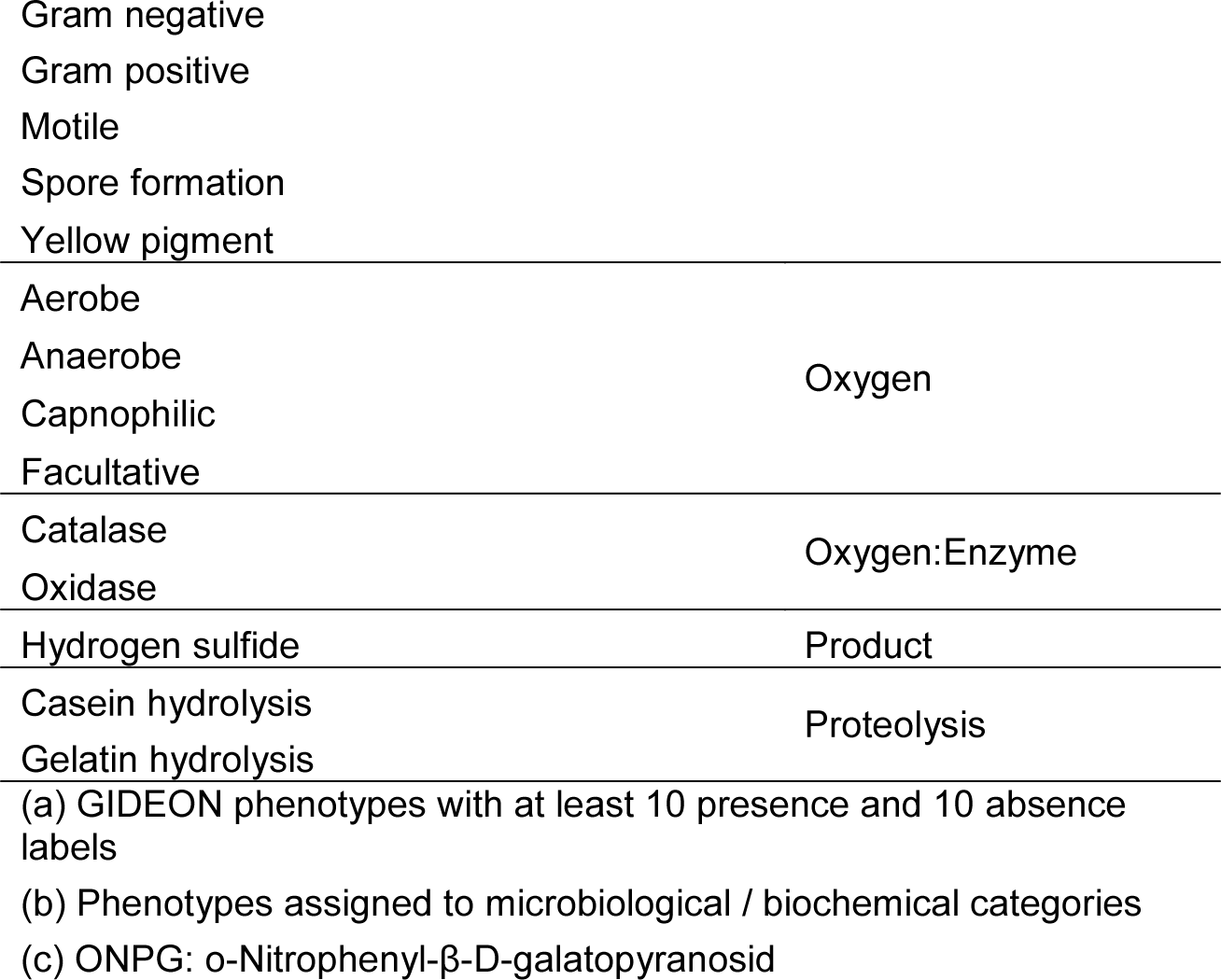

**Figure 2:**
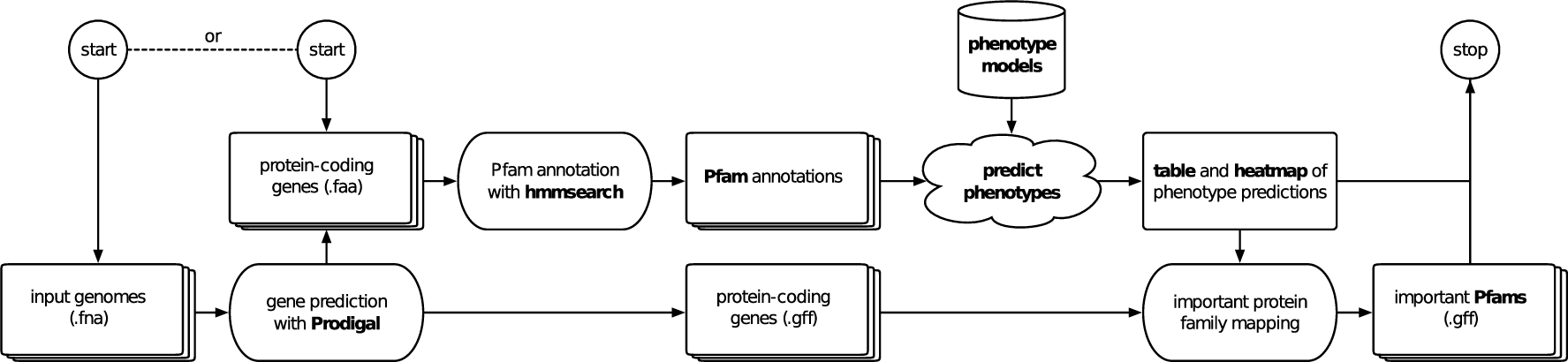
Work flow of Traitar. Input to the software can be genome sequene samples in nucleotide or amino acid FASTA format. Traitar predicts phenotypes based on pre-computed classification models and provides graphical and tabular output. In the case of nucleotide input, the protein families that are important for the phenotype predictions will be further mapped to the predicted protein-coding genes.

## Evaluation

We evaluated the two Traitar classifiers using ten-fold nested cross-validation on 234 bacterial species found in GIDEON (GIDEON I). The determined macro-accuracy (the accuracy balanced over all phenotypes) for the 67 GIDEON phenotypes was 82.6% for the phypat classifier and 85.5% for the phypat+PGL classifier; the accuracy (fraction of correct assignments averaged over all tested samples) for phypat was 88.1%, in comparison to 89.8% for phypat+PGL (Methods – Evaluation metrics; Table 2). Notably, Traitar classified 53 phenotypes with more than 80% macro-accuracy and 26 phenotypes with at least 90% macro-accuracy with one of the two classifiers (Figure 3, Supplementary Table 2). Phenotypes that could be predicted with very high confidence included the outcome of a ‘Methyl red’ test, ‘Spore formation’, oxygen requirement (i.e. ‘Anaerobe’ and ‘Aerobe’), ‘Growth on MacConkey agar’ or ‘Catalase’. Some phenotypes proved to be difficult to predict (60-70% macro-accuracy), which included ‘DNAse’, ‘myo-Inositol’ or ‘Yellow pigment’ and ‘Tartrate utilization’, regardless of which classifier was used. This might be caused by the relatively small number (<20) of positive (phenotype present) examples that were available.

**Table 2:**
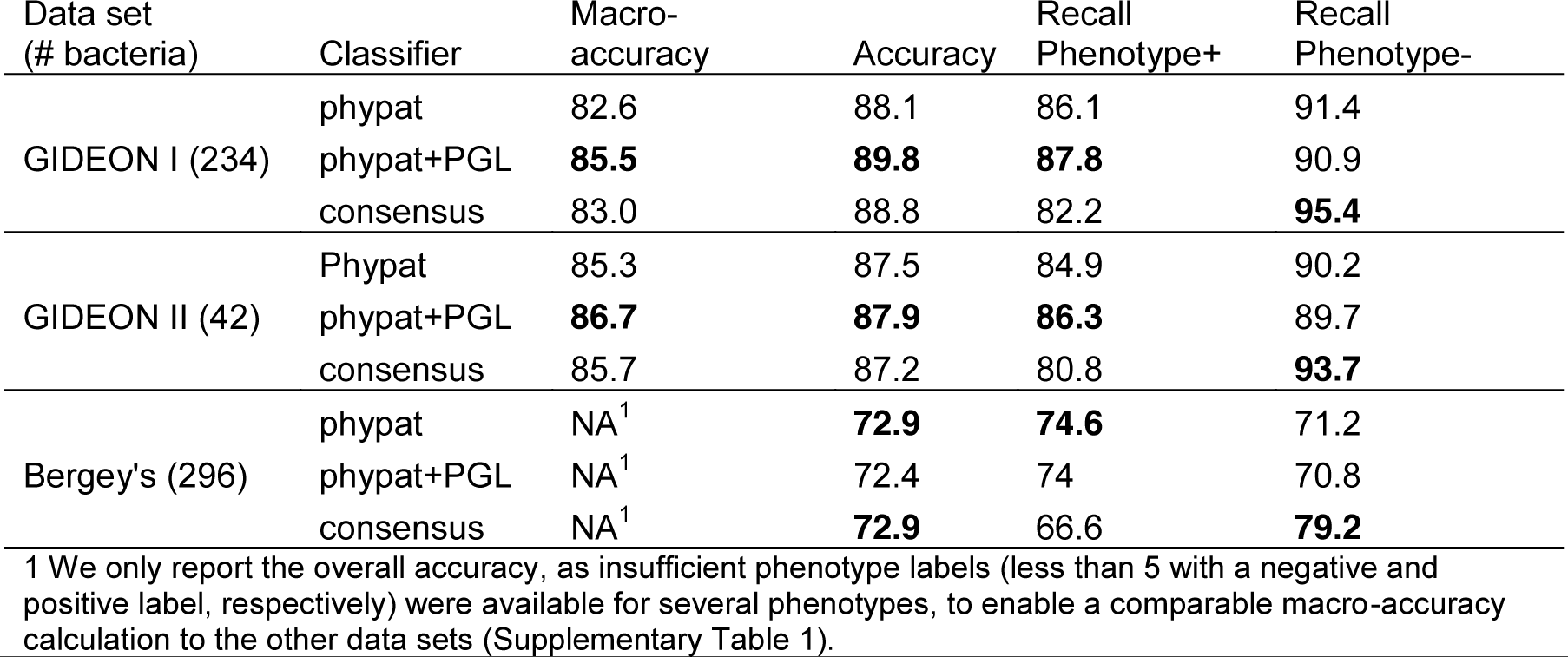
We evaluated the Traitar phypat and phypat+PGL phenotype classifiers and a consensus vote of both classifiers for 234 bacteria described in the Global Infectious Disease and Epidemiology Online Network (GIDEON) in a 10-fold nested cross-validation using different evaluation measures (Methods – Evaluation). Subsequently, we tested another 42 bacteria from GIDEON and 296 bacteria described in Bergey’s manual of systematic bacteriology for an independent performance assessment of the two classifiers.

| Data set<br>(# bacteria) | Classifier | Macro-<br>accuracy | Accuracy | Recall<br>Phenotype+ | Recall<br>Phenotype- |
| --- | --- | --- | --- | --- | --- |
| GIDEON I (234) | phypat | 82.6 | 88.1 | 86.1 | 91.4 |
|  | phypat+PGL | **85.5** | **89.8** | **87.8** | 90.9 |
|  | consensus | 83.0 | 88.8 | 82.2 | **95.4** |
| GIDEON II (42) | Phypat | 85.3 | 87.5 | 84.9 | 90.2 |
|  | phypat+PGL | **86.7** | **87.9** | **86.3** | 89.7 |
|  | consensus | 85.7 | 87.2 | 80.8 | **93.7** |
| Bergey’s (296) | phypat | NA<sup>1</sup> | **72.9** | **74.6** | 71.2 |
|  | phypat+PGL | NA<sup>1</sup> | 72.4 | 74 | 70.8 |
|  | consensus | NA<sup>1</sup> | **72.9** | 66.6 | **79.2** |

<sup>1</sup> We only report the overall accuracy, as insufficient phenotype labels (less than 5 with a negative and positive label, respectively) were available for several phenotypes, to enable a comparable macro-accuracy calculation to the other data sets (Supplementary Table 1).

**Figure 3:**
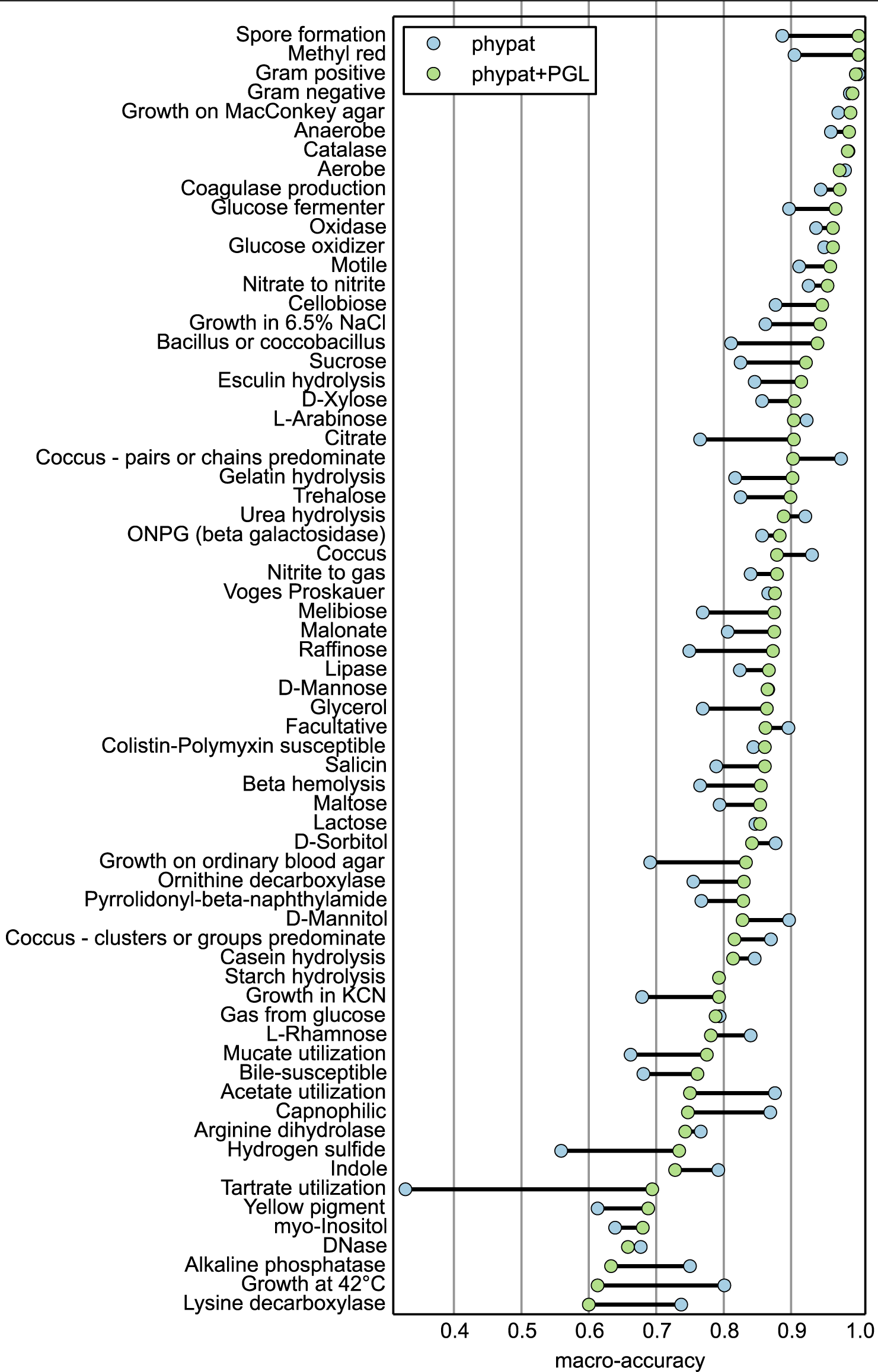
Macro-accuracy for each phenotype for the Traitar phypat and phypat+PGL phenotype classifiers determined in nested cross-validation on 234 bacterial species described in the Global Infectious Disease and Epidemiology Online Network (Methods – Evaluation metrics, Table 1, Supplementary Table 1).

For an independent assessment of Traitar’s classification performance we next tested Traitar on 42 bacterial species that had phenotype information available in GIDEON (GIDEON II), but were not used for learning the phenotype models (The Traitar software – Annotation). For calculation of the macro-accuracy, we considered only phenotypes represented by at least five phenotype-positive and five phenotype-negative bacteria. On these data, Traitar predicted the phenotypes with a macro-accuracy of 85.3% with the phypat classifier and 86.7% with the phypat+PGL classifier, and accuracies of 87.5% and 87.9%, respectively (Table 2). To investigate the performance of Traitar for bacterial genomes from a different data source, we next determined from two volumes of Bergey’s Manual of Systematic Bacteriology, namly ‘The Proteobacteria’ and ‘The Firmicutes’, the phenotypes of further sequenced bacteria that were not in our GIDEON I and II data sets (Supplementary Table 1, 4). In total, we thus identified phenotypes for another 296 sequenced bacterial species (The Traitar software – Annotation). Also for these bacteria, Traitar performed well but was less reliable than before, with accuracies for the phypat classifier of 72.9% and 72.1% for the phypat+PGL classifier (Table 2). This is likely due to the taxonomic differences of bacteria listed in GIDEON and Bergey’s and also because most of the bacteria in Bergey’s have only draft genomes available for phenotyping.

When combining the predictions of the phypat and phypat+PGL classifiers into a consensus vote, Traitar assigns phenotypes more reliably, while predicting less phenotype labels compared to the individuals classifiers (Table 2). Depending on the use case, Traitar can be used with performance characterized by different trade-offs between the recall of the phenotype-positive and the phenotype-negative classes.

## Performance per taxon at different ranks of the taxonomy

**Figure 4:**
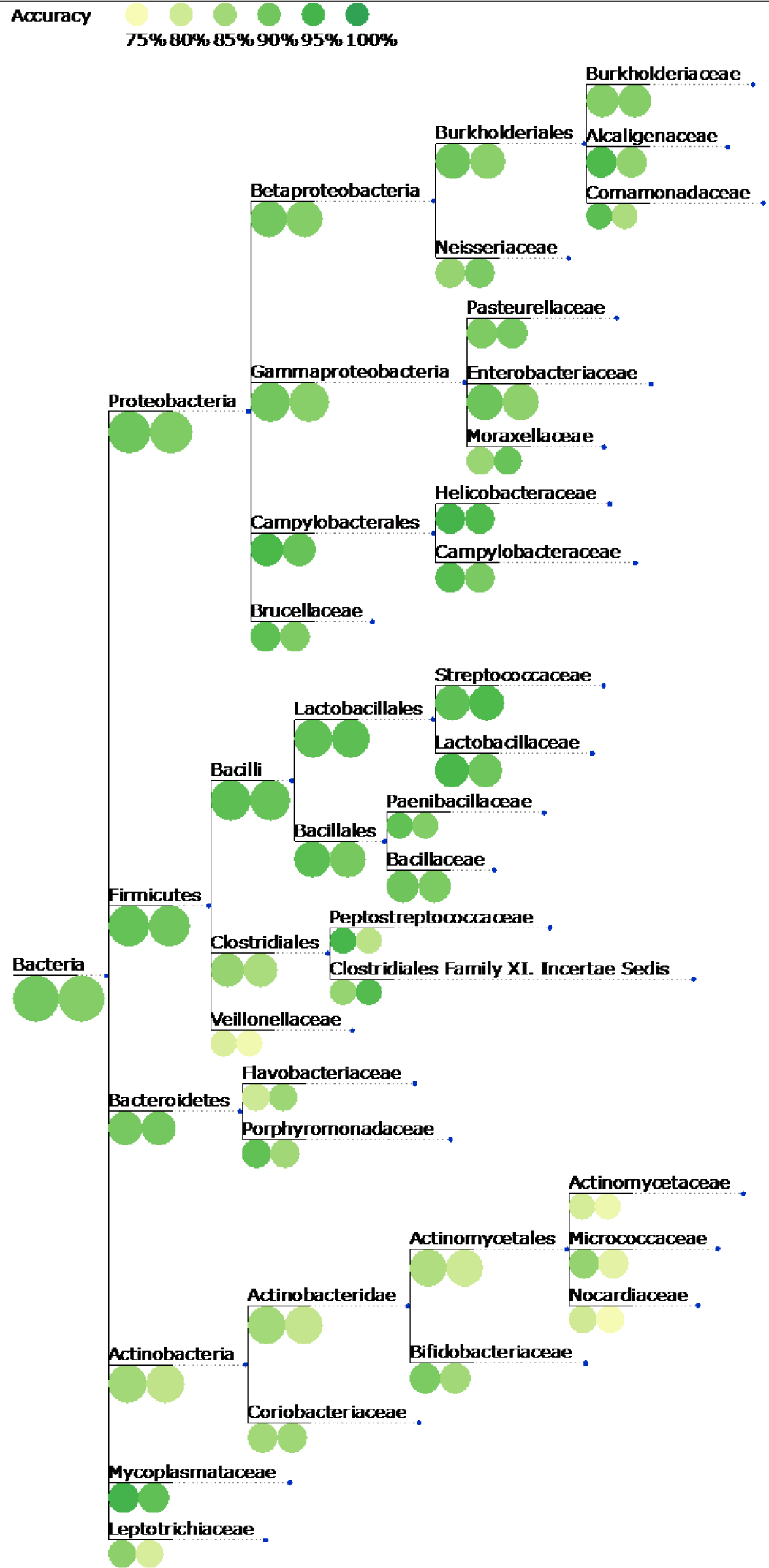
Classification accuracy for each taxon at different ranks of the NCBI taxonomy. For better visualization of names for the internal nodes, the taxon names are displayed on branches leading to the respective taxon node in the tree. The nested cross-validation accuracy obtained with Traitar for 234 bacterial species described in the Global Infectious Disease and Epidemiology Online Network was projected onto the NCBI taxonomy down to the family level. Colored circles at the tree nodes depict the performance of the phypat+PGL classifier (left-hand circles) and the phypat classifier (right-hand circles). The size of the circles reflects the number of species per taxon.

We investigated the performance of Traitar across the part of the bacterial tree of life represented in our data set. For this purpose, we evaluated the nested cross-validation performance of the phypat and phypat+PGL classifiers at different ranks of the NCBI taxonomy. For a given GIDEON taxon, we pooled all bacterial species that are descendants of this taxon. Figure 4 shows the accuracy estimates projected on the NCBI taxonomy from the domain level down to individual families. Notably, the accuracy of the phypat+PGL (phypat) classifier for the phyla covered by at least five bacterial species showed low variance and was high across all phyla, ranging from 84% (81%) for Actinobacteria over 90% (89%) for Bacteroidetes, 89% (90%) for Proteobacteria, 91% (90%) for Firmicutes to 91% (86%) for Tenericutes.

## Phenotyping incomplete genomes

GFMs or SAGs are often incomplete and thus we analyzed the effect of missing genome assembly parts onto the performance of Traitar. Rinke *et al.* used a single-cell sequencing approach to analyze poorly characterized parts of the bacterial and archaeal tree of life, the so-called ‘microbial dark matter’ (Rinke et al., 2013). They pooled 20 SAGs from the candidate phylum Cloacimonetes, formerly known as WWE1, to generate joint – more complete – genome assemblies that had at least a genome-wide average nucleotide identity of 97% and belonged to a single 16S-based operational taxonomic unit, namely *Cloacamonas acidaminovorans (Pelletier et al., 2008)*.

According to our predictions based on the joint assembly of the single-cell genomes, *C. acidaminovorans* is Gram-negative and is adapted to an anaerobic lifestyle, which agrees with the description by Rinke *et al.* (Figure 5). Traitar further predicted ‘Arginine dihydrolase’ activity, which is in line with the characterization of the species as an amino acid degrader (Rinke et al., 2013). Remarkably, the prediction of a bacillus or coco-bacillus shape agrees with the results of Limam *et al.* (Limam et al., 2014), who used a WWE1-specific probe and characterized the samples with fluorescence *in situ* hybridization. They furthermore report that members of the Cloacimonetes candidate phylum are implicated in anaerobic digestion of cellulose, primarily in early hydrolysis, which is in line with the very limited carbohydrate degradation spectrum found by Traitar.

**Figure 5:**
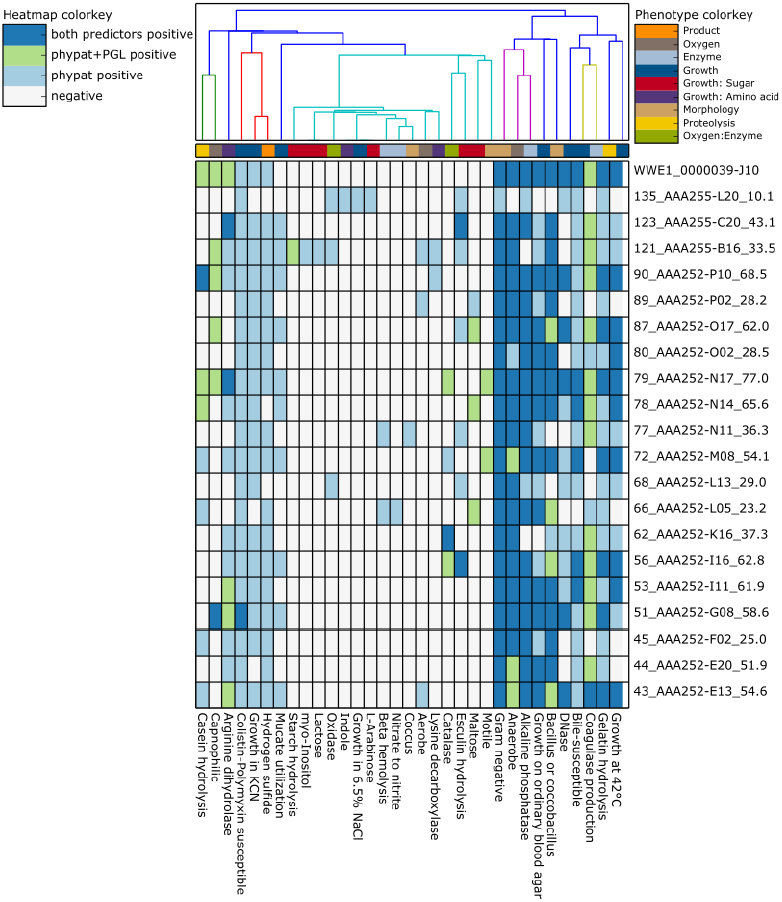
Single-cell phenotyping with Traitar. We used 20 genome assemblies with varying degrees of completeness from single cells of the Cloacimonetes candidate phylum and a joint assembly for phenotyping with Traitar. Shown is a heatmap of assembly samples vs. phenotypes, which is the standard visualization for phenotype predictions in Traitar. The origin of the phenotype’s prediction (Traitar phypat and/or Traitar phypat+PGL classifier) determines the color of the heatmap entries. The sample labels have their genome completeness estimates as suffixes. The colors of the dendrogram indicate similar phenotype distributions across samples, as determined by a hierarchical clustering with SciPy^1^.

Subsequently, we compared the predicted phenotypes for the SAGs to the predictions for the joint assembly. The phypat classifier recalled more of the phenotype predictions of the joint assembly based on the SAGs than the phypat+PGL classifier. However, the phypat+PGL classifier made fewer false positive predictions (Figure 6 a).

In the next experiment, we inferred phenotypes based on simulated GFMs, by subsampling from the coding sequences of each of the 42 bacterial genomes (GIDEON II). Starting with the complete set of coding sequences we randomly deleted genes from the genomes. For the obtained draft genomes with different degrees of completeness, we re-ran the Traitar classification and computed the accuracy measures, as before. We observed that the average fraction of phenotypes identified (macro-recall for the positive class) of the phypat+PGL classifier dropped more quickly with more missing coding sequences than that of the phypat classifier (Figure 6 b). However, at the same time, the recall of the negative class of the phypat+PGL classifier improved with a decreasing number of coding sequences, meaning that fewer but more reliable predictions were made.

Overall, the tradeoffs in the recall of the phenotype-positive and the phenotype-negative classes of the two classifiers resulted in a similar overall macro-accuracy across the range of tested genome completeness.

**Figure 6:**
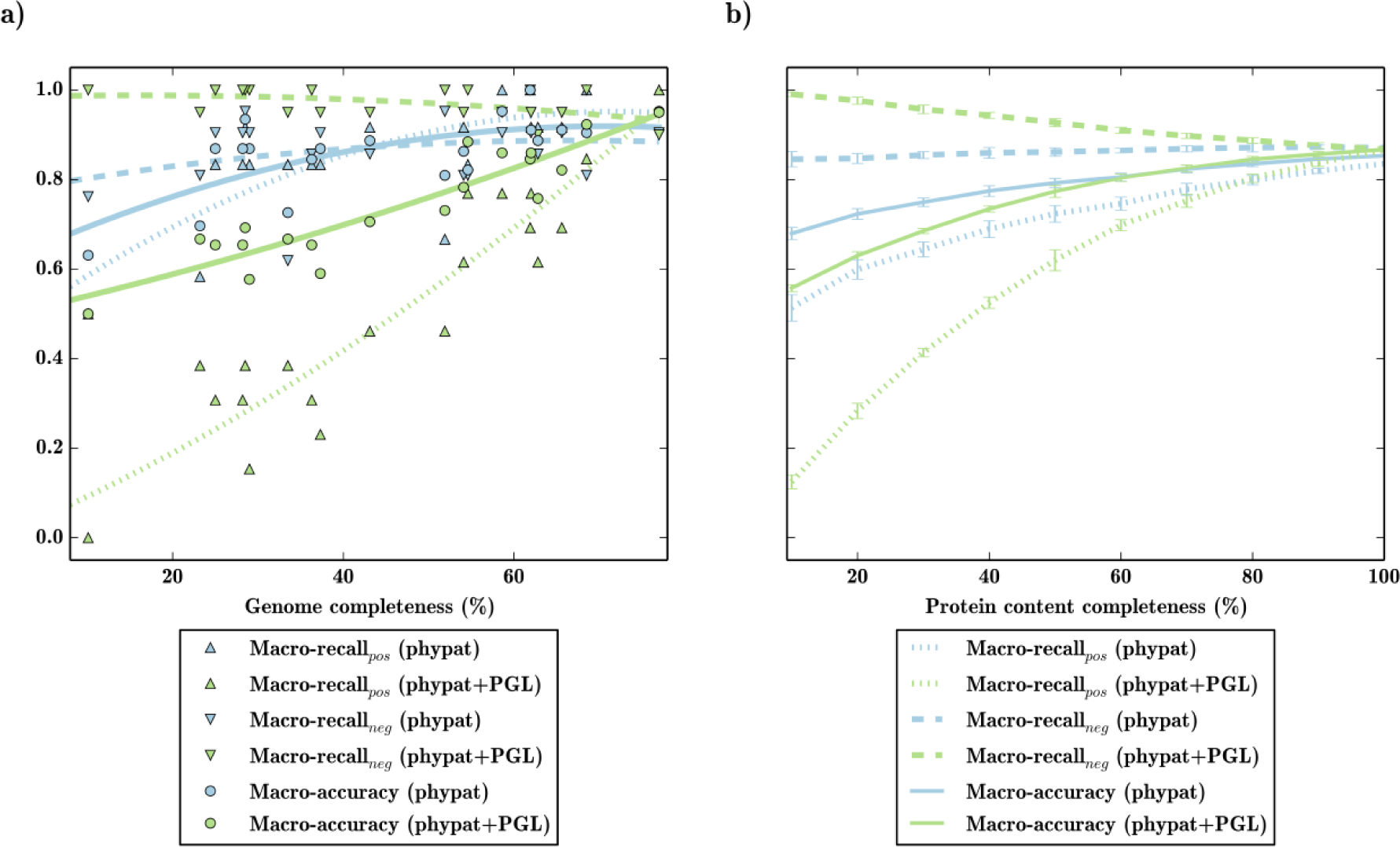
Phenotyping simulated draft genomes and single cell genomes. In (a) we used 20 genome assemblies with varying degrees of completeness from single cells of the Cloacimonetes candidate phylum and a joint assembly for phenotyping with the Traitar phypat and the Traitar phypat+PGL classifiers. Shown is the performance of the phenotype prediction vs. the genome completeness of the single cells with respect to the joint assembly. In (b) we simulated draft genomes based on an independent test set of 42 microbial (pan)genomes. The coding sequences of these genomes were down-sampled (10 replications per sampling point) and the resulting simulated draft genomes were used for phenotyping with the Traitar phypat and the Traitar phypat+PGL classifiers. We plotted various performance estimates (mean center values and and s.d. error bars shown) against the protein content completeness.

Thus, depending on the intended usage, a particular classifier can be chosen: we expect that the reliable predictions inferred with the phypat+PGL classifier and the more abundant, but less reliable predictions made with the phypat classifier will complement one another in different use cases for partial genomes recovered from metagenomic data.

By analyzing the protein families with assigned weights and the bias terms of the two classifiers, we found the phypat+PGL classifier to base its predictions primarily on the presence of protein families that were typical for the phenotypes. In contrast, the phypat classifier also took typically absent protein families from phenotype-positive genomes into account in its decision. More technically, the positive weights in models of the phypat classifier are balanced out by negative weights, whereas for the phypat+PGL classifier, they are balanced out by the bias term. By down-weighting the bias term for the phypat+PGL classifier by the protein content completeness, we could show that the accuracy of the phypat classifier could be increased over that of the phypat+PGL, regardless of the protein content completeness (data not shown). However, this requires knowledge of the protein content completeness for each genomic sample, which could be indirectly estimated using methods such as checkM (Parks et al., 2015).

## Traitar as a resource for gene target discovery

In addition to phenotype assignment, Traitar suggests the protein families relevant for the assignment of a phenotype (Methods – Majority feature selection, Table 3). We exemplarily demonstrate this capability here for three phenotypes that are already well-studied, namely ‘Motile’, ‘Nitrate to nitrite’ conversion and ‘L-arabinose’ metabolism. These phenotypes represent one each from the phenotype categories morphology, enzymatic activity and growth on sugar.

In general, we observed that the protein families important for classification can be seen to be gained and lost jointly with the respective phenotypes within the microbial phylogeny. Among the selected Pfam families that are important for classifying the motility phenotype were proteins of the flagellar apparatus and chemotaxis-related proteins (Table 3). Motility allows bacteria to colonize their preferred environmental niches. Genetically, it is mainly attributed to the flagellum, which is a molecular motor, and is closely related to chemotaxis, a process that lets bacteria sense chemicals in their surroundings. Motility also plays a role in bacterial pathogenicity, as it enables bacteria to establish and maintain an infection. For example, pathogens can use flagella to adhere to their host and they have been reported to be less virulent if they lack flagella (Josenhans and Suerbaum, 2002). Of 48 flagellar proteins described in (Liu and Ochman, 2007), four proteins (FliS, MotB, FlgD and FliJ) were sufficient for accurate classification of the motility phenotype and were selected by our classifier, as well as FlaE, which was not included in this collection. FliS (PF02561) is a known export chaperone that inhibits early polymerization of the flagellar filament FliC in the cytosol (Lam et al., 2010). MotB (PF13677), part of the membrane proton-channel complex, acts as the stator of the bacterial flagellar motor (Hosking et al., 2006). Traitar also identified further protein families related to chemotaxis, such as CZB (PF13682), a family of chemoreceptor zinc-binding domains found in many bacterial signal transduction proteins involved in chemotaxis and motility (Draper et al., 2011), and the P2 response regulator-binding domain (PF07194). The latter is connected to the chemotaxis kinase CheA and is thought to enhance the phosphorylation signal of the signaling complex (Dutta et al., 1999).

Nitrogen reduction in nitrate to nitrite conversion is an important step of the nitrogen cycle and has a major impact on agriculture and public health. Two types of nitrate reductases are found in bacteria: the membrane-bound Nar and the periplasmic Nap nitrate reductase (Moreno-Vivian et al., 1999), which we found both to be relevant for the classification of the phenotype: we identified all subunits of the Nar complex as being relevant for the ‘Nitrate to nitrite’ conversion phenotype (i.e. the gamma and delta subunit (PF02665, PF02613)), as well as Fer4_11 (PF13247), which is in the iron–sulfur center of the beta subunit of Nar. The delta subunit is involved in the assembly of the Nar complex and is essential for its stability, but probably is not directly part of it (Pantel et al., 1998). Traitar also identified the Molybdopterin oxidoreductase Fe4S4 domain (PF04879), which is bound to the alpha subunit of the nitrate reductase complex (Pantel et al., 1998). Traitor furthermore suggested NapB (PF03892) as relevant, which is a subunit of the periplasmic Nap protein and NapD (PF03927), which is an uncharacterized protein implicated in forming Nap (Moreno-Vivian et al., 1999).

**Figure 7:**
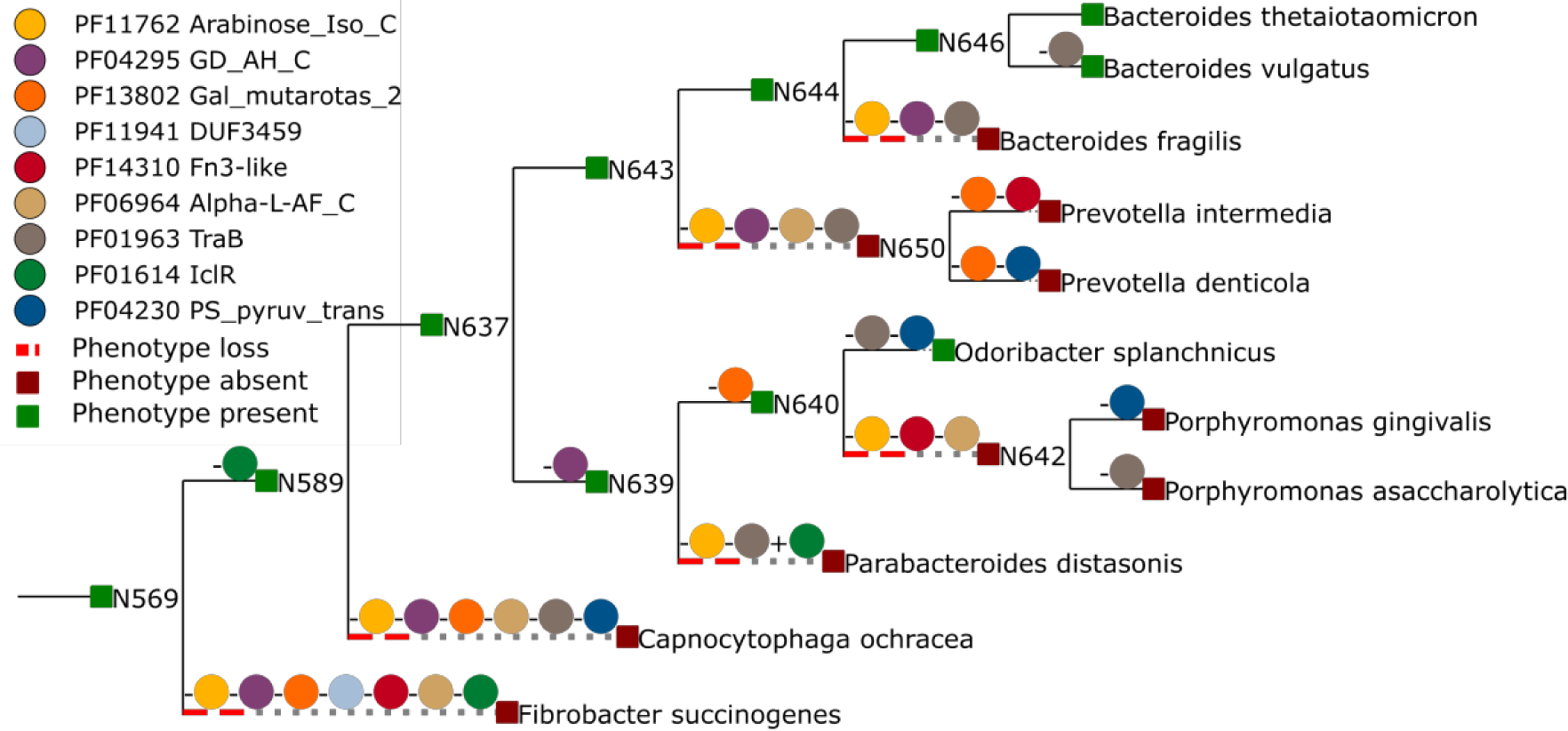
Phenotype gain and loss dynamics match protein family dynamics. We show the phenotype–protein family gain and loss dynamics for families identified as important by Traitar for the L-arabinose phenotype. Signed colored circles along the tree branches depict protein family gains (+) or losses (-). Taxon nodes are colored according to their inferred (ancestral) phenotype state.

**Table 3:**
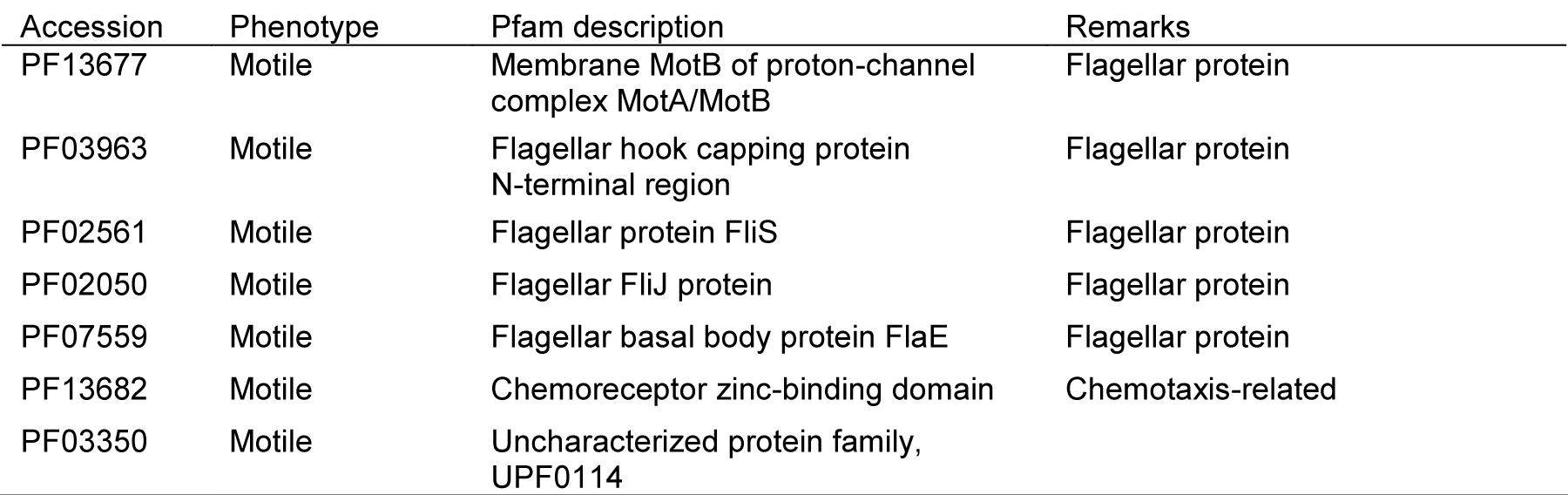
The most relevant Pfam families for classification of three important phenotypes: ‘Nitrate to Nitrite’, ‘Motility’ and ‘L-Arabinose’. We ranked the Pfam families with positive weights in the Traitar SVM classifiers by the correlation of the Pfam families with the respective phenotype labels across 234 bacteria described in the Global Infectious Disease and Epidemiology Online Network. Shown are the 10 highest ranking Pfam families along with their descriptions and a description of their phenotype-related function, where we found one.

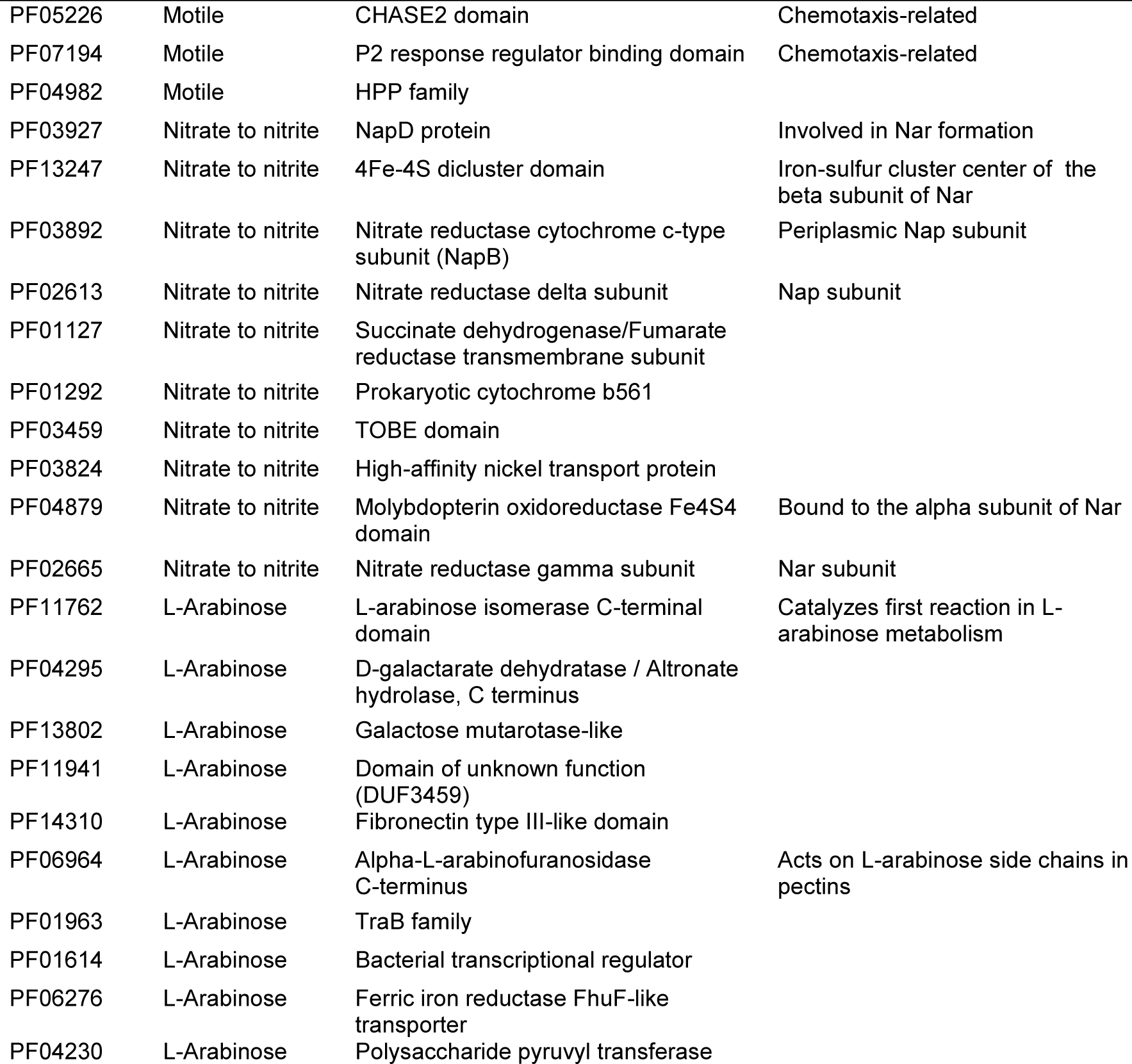

L-arabinose is major constituent of plant polysaccharides, which is located, for instance, in pectin side chains and is an important microbial carbon source (Martinez et al., 2008). Traitar identified the L-arabinose isomerase *C*-terminal domain (PF11762), which catalyzes the first step in L-arabinose metabolism – the conversion of L-arabinose into L-ribulose (Sa-Nogueira et al., 1997), as being important for realizing the L-arabinose metabolism. It furthermore suggested the *C*-terminal domain of Alpha-L-arabinofuranosidase (PF06964), which cleaves nonreducing terminal alpha-L-arabinofuranosidic linkages in L-arabinose-containing polysaccharides (Gilead and Shoham, 1995) and is also part of the well-studied L-arabinose operon in *Escherichia coli* (Sa-Nogueira et al., 1997).

## Phenotyping biogas reactor population genomes

We used Traitar to phenotype two novel Clostridiales species (unClos_1, unFirm_1) based on their genomic information reconstructed from metagenome samples. These were taken from a commercial biogas reactor operating with municipal waste (Frank et al., 2015). The genomes of unClos_1 and unFirm_1 were estimated to be 91% complete and 60% complete based on contigs ≥5 kb, respectively. Traitar predicted unClos_1 to utilize a broader spectrum of carbohydrates than unFirm_1 (Table 4). We cross-referenced our predictions with a metabolic reconstruction conducted by Frank *et al.* (under review; supplementary material). We considered all phenotype predictions that Traitar inferred with either the phypat or the phypat+PGL classifier. The manual reconstruction and predictions inferred with Traitar agreed to a great extent (Table 4). Traitar recalled 87.5% (6/7) of the phenotypes inferred via the metabolic reconstruction and also agreed to 81.8% (9/11) on the absent phenotypes. Notable exceptions were that Traitar only found a weak signal for ‘D-xylose’ utilization. A weak signal means that only a minority of the classifiers in the voting committee assigned these samples to the phenotype-positive class (Methods – Phenotype models). However, the metabolic reconstruction was also inconclusive with respect to xylose fermentation. Furthermore, Traitar only found a weak signal for ‘Glucose fermentation’ for unFirm_1. Whilst genomic analysis of unFirm_1 revealed the Embden–Meyerhof–Parnas (EMP) pathway, which would suggest glucose fermentation, gene-centric and metaproteomic analysis of this phylotype indicated that the EMP pathway was probably employed in an anabolic direction (gluconeogenesis); therefore unFirm_1 is also unlikely to ferment D-Mannose. This suggests that unFirm_1 is unlikely to ferment sugars and instead metabolizes acetate (also predicted by Traitar, Table 4) via a syntrophic interaction with hydrogen-utilizing methanogens.

Traitar predicted further phenotypes for both species that were not targeted by the manual reconstruction. One of these predictions was an anaerobic lifestyle, which is likely to be accurate, as the genomes were isolated from an anaerobic bioreactor environment. It also predicted them to be Gram-positive, which is probably correct, as the Gram-positive sortase protein family can be found in both genomes.

**Table 4.** Phenotype predictions for two novel Clostridiales species with genomes reconstructed from a commercial biogas reactor metagenome. Traitar output (yes, no, weak) was cross-referenced with phenotypes manually reconstructed based on Kyoto Encyclopedia of Genes and Genomes orthology annotation (Frank *et al.* submitted; supplementary material), which are primarily the fermentation phenotypes of various sugars. We considered all phenotype predictions that Traitar inferred with either the phypat or the phypat+PGL classifier. A weak prediction means that only a minority of the classifiers in the Traitar voting committee assigned this sample to the phenotype-positive class (Traitar phenotype). Table entries colored in red show a difference between the prediction and the reconstruction, whereas green denotes an overlap; yellow is inconclusive.

|  | unClos_1 | unFirm_1 |
| --- | --- | --- |
| Glucose | yes | weak |
| Acetate utilization | no | yes |
| Mannitol | yes | no |
| Starch hydrolysis | no | no |
| Xylose | weak | no |
| L-Arabinose | yes | no |
| Capnophilic | yes | no |
| Sucrose | yes | no |
| D-Mannose | yes | no |
| Maltose | yes | no |
| Arginine dihydrolase | no | yes |

This is a Gram-positive biomarker (Paterson and Mitchell, 2004). Furthermore, all Firmicutes known so far are Gram-positive (Goodfellow et al., 2012). Additionally, Traitar assigned ‘Motile’ and ‘Spore formation’ to unFirm_1, based on the presence of several flagellar proteins (e.g. FliM, MotB, FliS and FliJ) and the sporulation proteins CoatF and YunB.

## Discussion

We have developed Traitar, a software framework for predicting phenotypes from the protein family profiles of bacterial genomes. Traitar provides a quick and fully automated way of assigning 67 different phenotypes to bacteria based on the protein family content of their genomes.

Microbial trait prediction from phyletic patterns has been proposed in previous studies for a limited number of phenotypes (Feldbauer et al., 2015; Kastenmuller et al., 2009; Konietzny et al., 2014; Lingner et al., 2010; MacDonald and Beiko, 2010; Weimann et al., 2013). To our knowledge, the only currently available software for microbial genotype-phenotype inference is PICA, which is based on learning associations of clusters of orthologous genes (Tatusov et al., 2001) with traits (MacDonald and Beiko, 2010). Recently, PICA was extended by Feldbauer *et al.* for predicting eleven traits overall, optimized for large datasets and tested on incomplete genomes (Feldbauer et al., 2015). Traitar allows prediction of 67 phenotypes, including 60 entirely novel ones. It furthermore includes different prediction modes, one based on phyletic patterns, one additionally including a statistical model of protein family evolution for its predictions. Traitar also suggest associations between phenotypes and protein families. For three traits, we showed that several of these associations are to known key families of establishment of a particular trait, and that furthermore candidate families were suggested, that might serve as targets for experimental studies. Some of the phenotypes annotated in GIDEON are specific for the human habitat (such as ‘coagulase production’ or ‘growth on ordinary blood agar’) and the genetic underpinnings learned by Traitar could be interesting to study for infection disease research.

In cross-validation experiments with phenotype data from the GIDEON database, we showed that the Traitar phypat classifier has high accuracy in phenotyping bacterial samples. Considering ancestral protein family gains and losses in the classification, which is implemented in the Traitar phypat+PGL classifier, improves the accuracy compared to prediction from phyletic patterns only, both for individual phenotypes and overall. Barker *et al.* were first to note the phylogenetic dependence of genomic samples and how this can lead to biased conclusions (Barker and Pagel, 2005). MacDonald *et al.* selected protein families based on correlations with a phenotype and corrected for the taxonomy (MacDonald and Beiko, 2010). Here we accounted for the evolutionary history of the phenotype and the protein families in the classifier training itself to automatically improve phenotype assignment. We additionally demonstrated the reliability of the performance estimates by phenotyping, with a similar accuracy, an independent test dataset with bacteria described in GIDEON, which we did not use in the cross-validation. Traitar also reliably phenotyped a large and heterogenic collection of bacteria that we extracted from Bergey’s Manual of Systematic Bacteriology – mostly with only draft genomes available. We didn’t observe any bias towards specific taxa in GIDEON, but some of the phenotypes might be realized with different protein families in taxa that are less well represented indicated by the around 15% - 20% less reliable phenotyping results for bacteria described in Bergey’s manual of systematic bacteriology. We expect that the accuracy of the phenotype classification models already available in Traitar will further improve the more data will become available and can be incorporated into its training.

We found that Traitar can provide reliable insights into the metabolic capabilities of microbial community members even from partial genomes, which are very common for genomes recovered from single cells or metagenomes. One obvious limitation being for incomplete genomes, the absence of a phenotype prediction may be due to the absence of the relevant protein families from the input genomes. The analysis of both the SAGs and simulated genomes led us to the same conclusions: the phypat classifier is more suitable for exploratory analysis, as it assigned more phenotypes to incomplete genomes, at the price of more false positive predictions. In contrast, the phypat+PGL classifier assigned fewer phenotypes, but also made fewer false assignments. At the moment, genotype–phenotype inference with Traitar only takes into account the presence and absence of protein families of the bacteria analyzed. This information can be readily computed from the genomic and metagenomic data. Future research could focus also on integration of other ‘omics’ data to allow even more accurate phenotype assignments. Additionally, expert knowledge of the biochemical pathways that are used in manual metabolic reconstructions, for example, could be integrated as prior knowledge into the model in future studies.

For the phenotyping of novel microbial species, generating a detailed (manual) metabolic reconstruction such as the one by Frank *et al.* (submitted; supplementary material) is time-intensive. Furthermore, such reconstructions are usually focused on specific pathways and are dependent on the research question. This is not an option for studies with 10–50+ genomes, which are becoming more and more common in microbiology (Brown et al., 2015; Hess et al., 2011; Rinke et al., 2013). Traitar thus is likely to be particularly helpful for multi-genome studies. It furthermore may pick up on things outside of the original research focus and could serve as a seed or a first-pass method for a detailed metabolic reconstruction in future studies.

## Methods

### The Traitar software

In this section we first describe the Traitar annotation procedure. We proceed with the genome and phenotype data used for the training of Traitar phenotype models; afterwards we explain the training and illustrate how we considered ancestral protein family gains and losses in the models. Finally, we specify the requirements for running the Traitar software.

#### Annotation

In the case of nucleotide DNA sequence input, Traitar uses Prodigal (Hyatt et al., 2010) for gene prediction prior to Pfam family annotation. The amino acid sequences are then annotated in Traitar with protein families (Pfams) from the Pfam database (version 27.0) (Finn et al., 2014) using the hmmsearch command of HMMER 3.0 (Finn et al., 2011).

Each Pfam family has a hand-curated threshold for the bit score, which is set in such a way that no false positive is included (Punta et al., 2012). A fixed threshold of 25 is then applied to the bit score (the log-odds score) and all Pfam domain hits with an E-value above 10^-2^ are discarded. The resulting Pfam family counts (phyletic patterns) are turned into presence or absence values, as we found this representation to yield a favorable classification performance (Weimann et al., 2013).

#### Genome and phenotype data

We obtained our phenotype data from the GIDEON database (Berger, 2005). In GIDEON a bacterium is labeled either as phenotype-positive, -negative or strain-specific. In the latter case we discarded this phenotype label. The GIDEON traits can be grouped into the categories the use of various substrates as source of carbon and energy for growth, oxygen requirement, morphology, antibiotic susceptibility and enzymatic activity (Table 1, Supplementary Table 1). We only considered phenotypes that were available in GIDEON for at least 20 bacteria, with a minimum of 10 bacteria annotated as positive (phenotype presence) for a given phenotype and 10 as negative (phenotype absence) to enable a robust and reliable analysis of the respective phenotypes. Furthermore, to be included in the analysis, we required each bacterial sample to have:

a) at least one annotated phenotype,
b) at least one sequenced strain,
c) a representative in the sTOL.

In total, we extracted 234 species-level bacterial samples with 67 phenotypes with sufficient total, positive and negative labels from GIDEON (GIDEON I). GIDEON associates these bacteria with 9305 individual phenotype labels, 2971 being positive and 6334 negative (Supplementary Table 1, 3). GIDEON species that had at least one sequenced strain available but were not part of the sTOL tree were set aside for a later independent assessment of the classification accuracy. In total, this additional dataset comprised further 42 unique species with 58 corresponding sequenced bacterial strains (GIDEON II, Supplementary Table 1, 4). We obtained 1836 additional phenotype labels for these bacteria, consisting of 574 positive and 1262 negative ones. We searched the Firmicutes and Proteobacteria volumes of Bergey’s systematic bacteriology specifically for further bacteria not represented so far in the GIDEON data sets (Goodfellow et al., 2012). In total, we obtained phenotype data from Bergey’s for 206 Firmicutes and 90 Proteobacteria with a total of 1152 positive labels and 1376 negative labels (Supplementary Table 1, 5). As in GIDEON, in Bergey’s the phenotype information is usually given on the species level.

We downloaded the coding sequences of all complete bacterial genomes that were available via the NCBI FTP server under ftp://ftp.ncbi.nlm.nih.gov/genomes/ as of 11 May 2014 and genomes from the PATRIC data base as of September 2015 (Wattam et al., 2014). These were annotated with Traitar. For bacteria with more than one sequenced strain available, we chose the union of the Pfam family annotation of the single genomes to represent the pangenome Pfam family annotation, as in (Liu et al., 2006).

#### Phenotype models

We represented each phenotype from the set of GIDEON phenotypes across all genomes as a vector ***yp***, and solved a binary classification problem using the matrix of Pfam phyletic patterns *XP* across all genomes as input features and ***yp*** *a*s the binary target variable (Supplementary Figure 1). For classification, we relied on support vector machines (SVMs), which are a well-established machine learning method (Boser et al., 1992). Specifically, we used a linear L1-regularized L2-loss SVM for classification as implemented in the LIBLINEAR library (Fan et al., 2008). For many datasets, linear SVMs achieve comparable accuracy to SVMs with a nonlinear kernel but allow faster training. The weight vector of the separating hyperplane provides a direct link to the Pfam families that are relevant for the classification. L1-regularization enables feature selection, which is useful when applied to highly correlated and high-dimensional datasets, as used in this study (Zou and Hastie, 2005). We used the interface to LIBLINEAR implemented in scikit-learn (Pedregosa et al., 2011). For classification of unseen data points – genomes without available phenotype labels supplied by the user – Traitar uses a voting committee of five SVMs with the best single cross-validation accuracy (Methods – Nested cross-validation). Traitar then assigns each unseen data point to the majority class (phenotype presence or absence class) of the voting committee.

#### Ancestral protein family and phenotype gains and losses

We constructed an extended classification problem by including ancestral protein family gains and losses, as well as the ancestral phenotype gains and losses in our analysis, as implemented in GLOOME (Cohen and Pupko, 2011). Barker *et al.* report that common methods for inferring functional links between genes, that do not take the phylogeny into account, suffer from high rates of false positives (Barker and Pagel, 2005). Here, we jointly derived the classification models from the observable phyletic patterns and phenotype labels, and from phylogenetically unbiased ancestral protein family and phenotype gains and losses, that we inferred via a maximum likelihood approach from the observable phyletic patterns on a phylogenetic tree, showing the relationships among the samples. (Supplementary Figure 1). Ancestral character state evolution in GLOOME is modeled via a continuous-time Markov process with exponential waiting times. The gain and loss rates are sampled from two independent gamma distributions (Cohen and Pupko, 2010).

GLOOME needs a binary phylogenetic tree with branch lengths as input. The taxonomy of the National Center for Biointechnology Information (NCBI) and other taxonomies are not suitable, because they provide no branch length information. We used the sequenced tree of life (sTOL) (Fang et al., 2013), which is bifurcating and was inferred with a maximum likelihood approach based on unbiased sampling of structural protein domains from whole genomes of all sequenced organisms (Gough et al., 2001). We employed GLOOME with standard settings to infer posterior probabilities for the phenotype and Pfam family gains and losses from the Pfam phyletic patterns of all NCBI bacteria represented in the sTOL and the GIDEON phenotypes. Each GIDEON phenotype *p* is available for a varying number of bacteria. Therefore, for each phenotype, we pruned the sTOL to those bacteria that were both present in the NCBI database and had a label for the respective phenotype in GIDEON. The posterior probabilities of ancestral Pfam gains and losses were then mapped onto this GIDEON phenotype-specific tree (Gps-sTOL, Supplementary Figure 2).

Let *B* be the set of all branches in the sTOL and *P* be the set of all Pfam families. We then denote the posterior probability *g*_*ij*_ of an event *a* for a Pfam family *pf* to be a gain event on branch *b* in the sTOL computed with GLOOME as:

*g_ij_ = P(a = gain|i = b, j = pf) ∀ i ∈ B, ∀ j ∈ P,*

and the posterior probability of *a* to be a loss event for a Pfam family *p* on branch *b* as:

*l_ij_ = P(a = loss|i = b, j = pf) ∀ i ∈ B, ∀ j ∈ P.*

We established a mapping *f*:*B*′ → *B* between the branches of the sTOL *B* and the set of branches *B*′ of the Gps-sTOL (Supplementary Figure 2). This was achieved by traversing the tree from the leaves to the root.

There are two different scenarios for a branch *b*′ in *B*′ to map to the branches in B:

a) Branch *b*′ in the Gps-sTOL derives from a single branch b in the sTOL: *f*(*b*′) = {*b*}. The posterior probability of a Pfam gain inferred in the Gps-sTOL on branch *b*′ consequently is the same as that on branch in the sTOL *g_bij_* = *g_bj_∀ j∈P*.
b) Branch *b*′ in the Gps-sTOL derives from *m* branches *b*_1_, …, *b*_*m*_ in the sTOL: *f*(*b*′) = {*b*_1_, …, *b*_*m*_} (Supplementary figure 2). In this case, we iteratively calculated the posterior probabilities for at least one Pfam gain *g*′ on branch *b*′ from the posterior probabilities for a gain *g*′_*b*_1_*j*_ from the posterior probabilities *g*_1_, …, *g*_*m*_ of a gain on branches *b*_1_, …, *b*_*m*_ with the help of h:

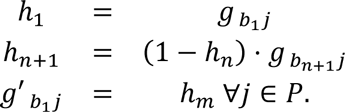

Inferring the Gps-sTOL Pfam posterior loss probabilities *l*′_*ij*_ from the sTOL posterior Pfam loss probabilities is analogous to deriving the gain probabilities. The posterior probability for a phenotype *p* to be gained *g*_*ip*_′ or lost *l*_*ip*_′ can be directly defined for the Gps-sTOL in the same way as for the Pfam probabilities.

For classification, we did not distinguish between phenotype or Pfam gains or losses, assuming that the same set of protein families gained with a phenotype will also be lost with the phenotype. This assumption simplified the classification problem. Specifically, we proceeded in the following way:

1. We computed the joint probability *x*_*ij*_ of a Pfam family gain or loss on branch *b*′ and the joint probability *y*_*j*_ of a phenotype gain or loss on branch *b*′:

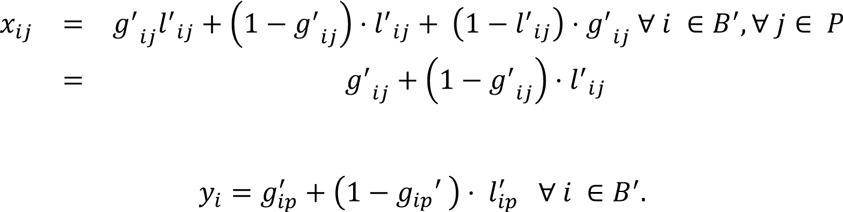
2. Let ***x***_***i***_ be a vector representing the probabilities *x*_*ij*_ for all Pfam families *j* ∈ *P* on branch *b*_*i*_. We discarded any samples (***x***_*i*_, *y*_*i*_) that had a probability for a phenotype gain or loss *y*_*i*_ above the reporting threshold of GLOOME but below a threshold *t*. We set the threshold *t* to 0.5. This defines the matrix X and the vector **y** as:

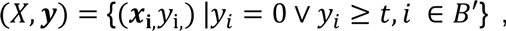 By this means, we avoided presenting the classifier with samples corresponding to uncertain phenotype gain or loss events and used only confident labels in the subsequent classifier training instead.
3. We inferred discrete phenotype labels ***y***′ by applying this threshold *t* to the joint probability *y*_*i*_ for a phenotype gain or loss to set up a well-defined classification problem with a binary target variable. Whenever the probability for a phenotype to be gained or lost on a specific branch was larger than *t*, the event was considered to have happened:

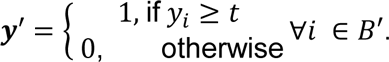
4. Finally, we formulated a joint binary classification problem for each target phenotype ***yp*** and the corresponding gain and loss events ***y***′ the phyletic patterns *XP*, and the Pfam gain and loss events *X*, which we solved again with a linear L1-regularized L2-loss SVM. We applied this procedure for all GIDEON phenotypes under investigation.

#### Software Requirements

Traitar can be run on a standard laptop with Linux/Unix. The runtime (wallclock time) for annotating and phenotyping a typical microbial genome with 3 Mbp is 9 minutes (3 min/Mbp) on an Intel(R) Core(TM) i5-2410M dual core processor with 2.30 GHz, requiring only a few megabytes of memory.

## Cross-validation

We employed cross-validation to assess the performance of the classifiers individually for each phenotype. For a given phenotype, we divided the bacterial samples that were annotated with that phenotype into ten folds. Each fold was selected once for testing the model, which was trained on the remaining folds. The optimal regularization parameter *C* needed to be determined independently in each step of the cross-validation; therefore, we employed a further inner cross-validation using the following range of values for the parameter *C*: 10^−3^, 10^−2^ · 0.7, 10^−2^ · 0.5, 10^−2^ · 0.2, 10^−2^ · 0.1, …, 1. In other words, for each fold kept out for testing in the outer cross-validation, we determined the value of the parameter *C* that gave the best accuracy in an additional tenfold cross-validation on the remaining folds. This value was then used to train the SVM model in the current outer cross-validation step. Whenever we proceeded to a new cross-validation fold, we re-computed the ancestral character state reconstruction of the phenotype with only the training samples included (Ancestral protein family and phenotype gains and losses). This procedure is known as nested cross-validation (Ruschhaupt et al., 2004).

The bacterial samples in the training folds imply a Gps-sTOL in each step of the inner and outer cross-validation without the samples in the test fold. We used the same procedure as before to map the Pfam gains and losses inferred previously on the Gps-sTOL onto the tree defined by the current cross-validation training folds. Importantly, the test error is only estimated on the observed phenotype labels rather than on the inferred phenotype gains and losses.

## Evaluation metrics

We used evaluation metrics from multi-label classification theory for performance evaluation (Manning et al., 2008). We determined the performance for the individual phenotype-positive and the phenotype-negative classes based on the confusion matrix of true positive (*TP*), true negative (*TN*), false negative (*FN*) and false positive (*FP*) samples from their binary classification equivalents by averaging over all phenotypes. We utilized two different accuracy measures for assessing multi-class classification performance (i.e. the accuracy pooled over all classification decisions and the macro-accuracy). Macro-accuracy represents an average over the accuracy of the individual binary classification problems and we computed this from the macro-recall of the phenotype-positive and the phenotype-negative classes as follows:

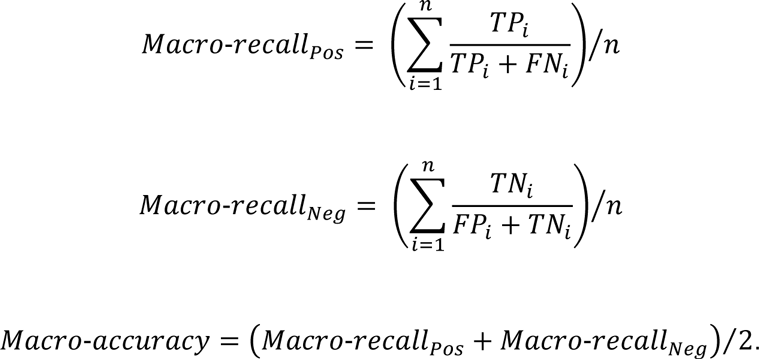

However, if there are only few available labels for some phenotypes, the variance of the macro-accuracy will be high and this measure cannot be reliably computed anymore; it cannot be computed at all if no labels are available. The accuracy only assesses the overall classification performance without consideration of the information about specific phenotypes. Large classes dominate small classes (Manning et al., 2008).

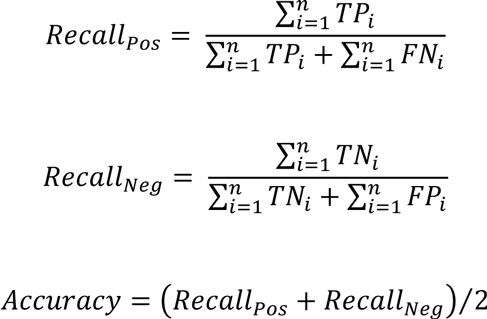

## Majority feature selection

The weights in linear SVMs can directly be linked to features that are relevant for the classification. We identified the most important protein families used as features from the voting committee of SVMs consisting of the five most accurate models, which were also used for classifying new samples. If the majority, which is at least three predictors, included a positive value for a given protein family, we added this feature to the list of important features. We further ranked these protein families features by their correlation with the phenotype using Pearson’s correlation coefficient.

## Acknowledgements

We thank Andreas Klötgen, David Lähnemann, Susanne Reimering and Alexander 10 Sczyrba for providing helpful comments on the manuscript; Johannes Dröge and Jens Loers for reviewing the Traitar software and Gary Robertson for helping to set up the Traitar web service. JAF and PBP are supported by a grant from the European Research Council (336355-MicroDE).

## Footnotes

1 http://docs.scipy.org/doc/scipy/reference/cluster.hierarchy.html

---

**Supplementary Figure 1.**
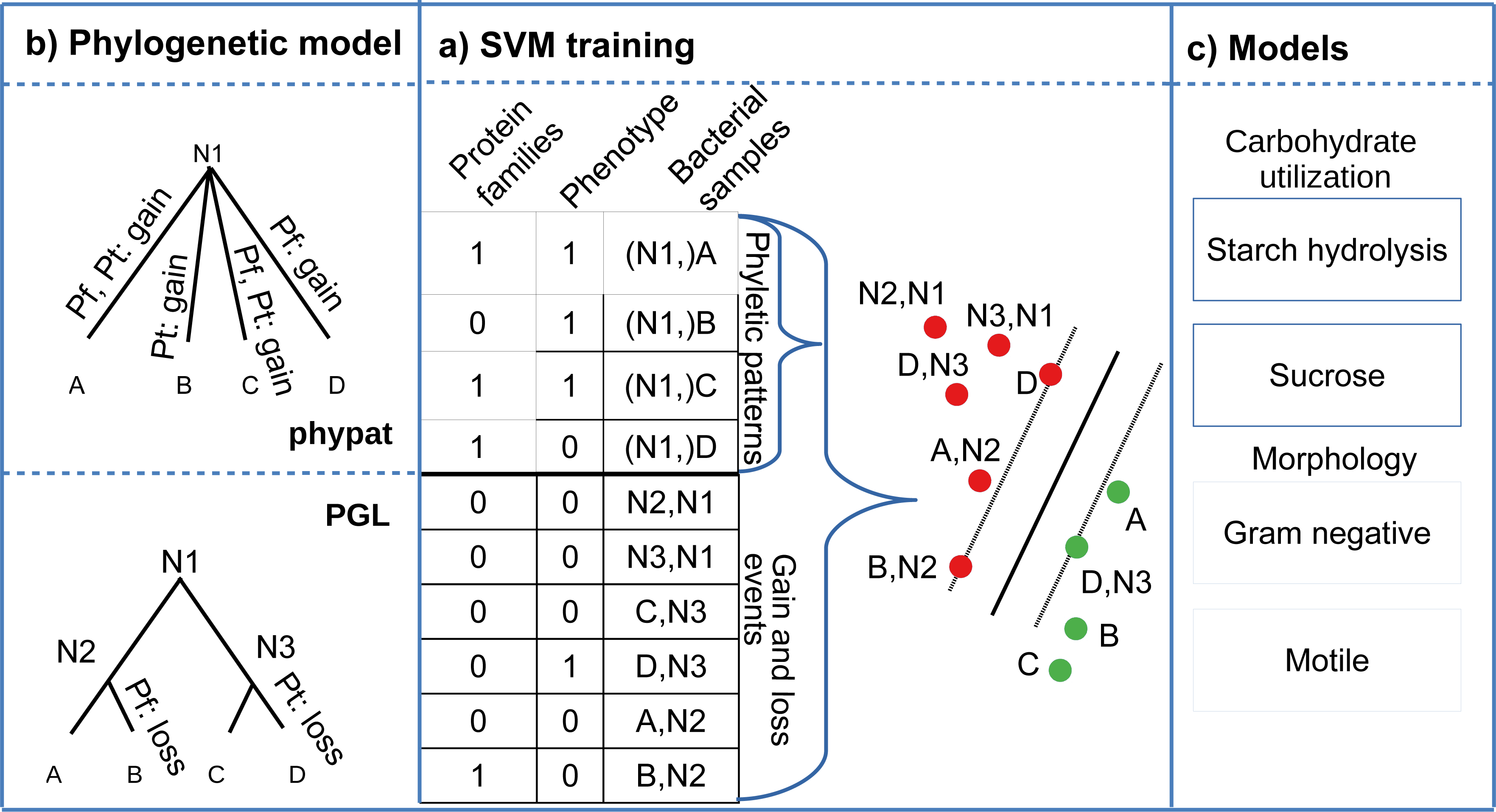
Schematic overview of the Traitar phenotype model training. (a) The phenotype and Pfam protein family phyletic patterns correspond to gain events on a star-shaped phylogenetic tree. Alternatively, we reconstructed the ancestral Pfam family and phenotype gain and loss events on the sequenced Tree of Life. (b) We trained a support vector machine classifier either on the phyletic patterns and on the ancestral gain and loss events, or solely on the phyletic patterns. (c) In this way, we inferred classification models for all available phenotypes.

**Supplementary Figure 1.**
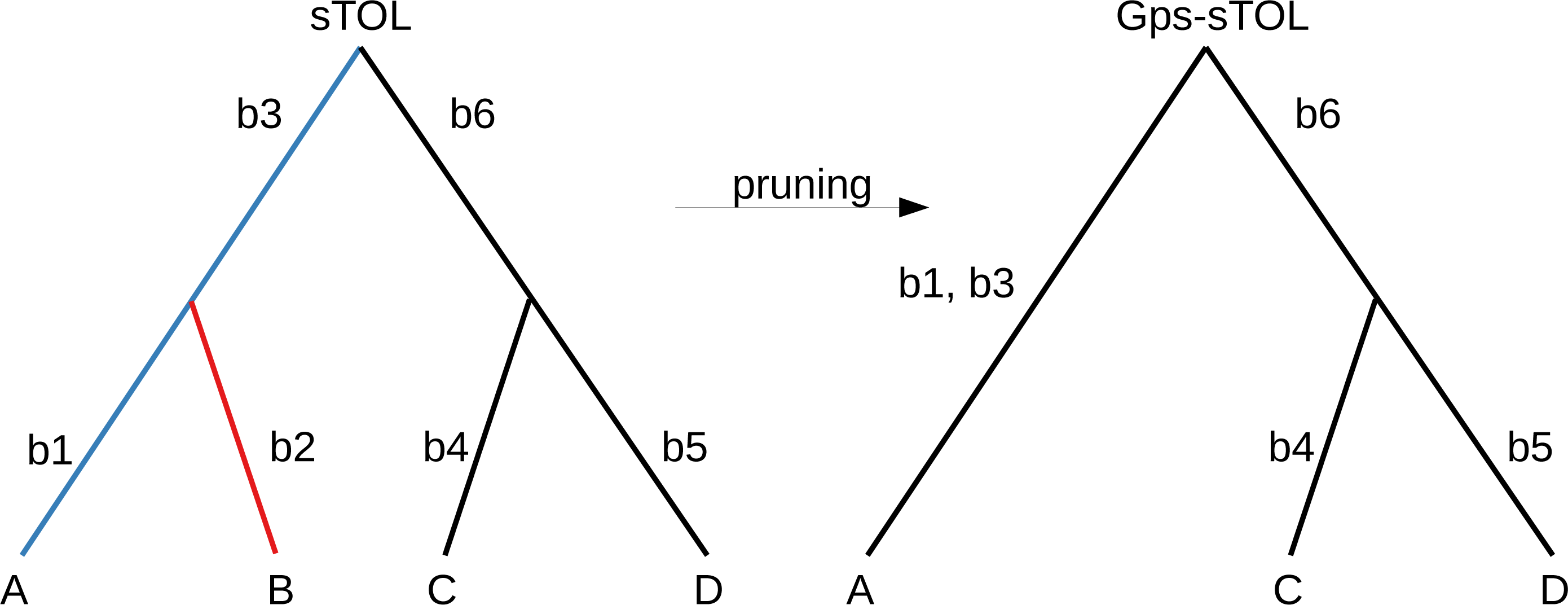
Sequenced Tree of Life (sTOL) and GIDEON phenotype-specific Tree of Life (Gps-sTOL) correspondence. The phenotype label for Sample B is not available. Consequently, only branches b_4_, b_5_ and b_6_are also found in the Gps-sTOL. The posterior probabilities for a Pfam gain or loss are the same for b_4_, b_5_ and b_6_ in both trees. Branches b_1_ and b_3_ (blue) are collapsed into a single branch. The posterior probability for a gain on branch b_1,3_, g_b1,3_ is computed from the posterior probability for a Pfam gain for b_1_and b_3_ as follows: g_b1,3_ = g_b1_ + (1 – g_b1_). g_b3_ Branch b_2_(red) in the sTOL does not have an analog in the Gps-sTOL.

